# Ecosystem photosynthesis in land-surface models: a first-principles approach

**DOI:** 10.1101/2021.05.07.442894

**Authors:** Giulia Mengoli, Anna Agustí-Panareda, Souhail Boussetta, Sandy P. Harrison, Carlo Trotta, I. Colin Prentice

## Abstract

Vegetation regulates land-atmosphere water and energy exchanges and is an essential component of land-surface models (LSMs). However, LSMs have been handicapped by assumptions that equate acclimated photosynthetic responses to environment with fast responses observable in the laboratory. These time scales can be distinguished by including specific representations of acclimation, but at the cost of further increasing parameter requirements. Here we develop an alternative approach based on optimality principles that predict the acclimation of carboxylation and electron-transport capacities, and a variable controlling the response of leaf-level carbon dioxide drawdown to vapour pressure deficit (VPD), to variations in growth conditions on a weekly to monthly time scale. In the “P model”, an optimality-based light-use efficiency model for gross primary production (GPP) on this time scale, these acclimated responses are implicit. Here they are made explicit, allowing fast and slow response time-scales to be separated and GPP to be simulated at sub-daily timesteps. The resulting model mimics diurnal cycles of GPP recorded by eddy-covariance flux towers in a temperate grassland and boreal, temperate and tropical forests, with no parameter changes between biomes. Best performance is achieved when biochemical capacities are adjusted to match recent midday conditions. This model suggests a simple and parameter-sparse method to include both instantaneous and acclimated responses within an LSM framework, with many potential applications in weather, climate and carbon - cycle modelling.

**Plain Language Summary:** Vegetation regulates the exchanges of energy, water and carbon dioxide between the land and the atmosphere. Numerical climate models represent these processes, focusing mainly on their rapid variations in response to changes in the environment (including temperature and light) on timescales of seconds to hours. However, plants also adjust their physiology to environmental changes over longer periods within the season. Here we have adapted a simple model that formulates plant behaviour in terms of optimal trade-offs between different processes, so it simulates processes on both time scales. This model correctly reproduces the daily cycle of carbon dioxide uptake by plants, as recorded in different kinds of vegetation. We show that plants optimize their behaviour for midday conditions, when the light is greatest, and adjust to longer-term environmental variations on a timescale of about a week to a month. The model conveniently avoids the need to give specific, fixed values to physiological variables (such as photosynthetic capacity) for different types of plants. The optimality assumptions mean that the model gives equally good results in tropical, temperate and boreal forests, and in grasslands, using the same equations, and a very small number of input variables that are constant across the world.

**Key Points:**

- Optimality theory is used to develop a simple model incorporating fast and acclimated responses of photosynthesis and stomatal conductance
- Biogeochemical photosynthetic capacities adjust to midday light conditions
- The new model simulates gross primary production on sub-daily timesteps across a range of different vegetation types and climate

## 1 Introduction

Vegetation plays a key role in the Earth system, regulating carbon, water and energy exchanges between vegetation and atmosphere. Transpiration, photosynthesis and respiration are the main processes that govern these exchanges and link vegetation to the climate (Bonan et al., 1992, 2003, 2008). Evapotranspiration (ET, with accompanying latent-heat release) and photosynthesis are tightly linked. Transpiration is the dominant component of ET, hence plants are the main conduit of water from the soil to the atmosphere. Plants control both transpiration and the flux of carbon dioxide (CO_2_) into leaves by regulating the opening or closing of stomata.

Through photosynthesis plants convert solar radiation into growth, storing carbon that otherwise would remain in the atmosphere as a climate-modifying greenhouse gas. CO_2_ is removed from the atmosphere by photosynthesis, but released again by autotrophic (plant) and heterotrophic (soil microbial) respiration (Ciais et al., 2013). Contemporary land surface models (LSMs) represent all these interactions.

Plants respond to environmental changes on different timescales. Fast (instantaneous) responses occur on timescales from seconds to hours; these are the plant responses to environmental stimuli before any type of physiological, structural or biochemical adjustment occurs. Longer-term responses (acclimation) occur over time scales of days to weeks (Mäkelä et al., 2019) or longer (Prentice and Cowling, 2013; Smith and Dukes, 2013). Plant acclimation is manifested as alterations in the short-term response functions of physiological processes (Smith and Dukes, 2013). Key photosynthetic traits, such as the maximum rate of carboxylation (*V*_cmax_) or the maximum rate of electron transport (*J*_max_) vary systematically with growth conditions, both in space and in time (Rogers et al., 2017; Togashi et al., 2018).

Originally LSMs used prescribed, plant functional type (PFT)-dependent values for photosynthetic traits. Modern versions of these models recognise the spatial and temporal variability of these traits within PFTs as a function of environmental conditions and thus include dynamic responses of photosynthetic (e.g. ORCHIDEE, JSBACH; see Fig.3 and Table 3 in Smith and Dukes, 2013) and (autotrophic) respiratory processes to temperature (e.g. JULES, CLM4.5, Atkin et al., 2008; Atkin et al., 2015; Lombardozzi et al., 2015; Kumarathunge et al., 2019). The approach used to account for plant acclimation remains a model parametrization, and therefore the differences between PFTs are maintained (Kattge & Knorr, 2007; Atkin 2008; Lawrence et al., 2019). Furthermore, the inclusion of acclimation generally involves additional parameters, with a consequent increase in model complexity (Fisher and Koven, 2020). Attempts have been made to include plant acclimation to light in soil-vegetation-atmosphere (SVAT) models (e.g. Meir et al., 2002) and terrestrial biosphere (TBM) models (e.g. Luo and Keenan et al., 2020) but most current LSMs do not address all aspects of acclimation. Many studies (e.g. Walker et al., 2017; Smith et al., 2019) have stressed the importance of including acclimation in models — using photosynthetic parameters that vary according to the climate — and indicated that this should lead to improved future projections. It has also been suggested that models that do not account for acclimation might overestimate the positive feedback between climate and vegetation in future scenarios (Smith et al., 2017).

Many current LSMs however continue to ignore acclimation and so, by considering only the instantaneous responses, they make no distinction between fast and slow processes. LSMs make inconsistent future projections of changes in the carbon and water cycles under the same future scenarios (Ciais et al., 2013; Prentice et al., 2015). They do not predict global primary production and its interannual variability correctly (Anav et al., 2015), and differ greatly, for example, in the responses of photosynthesis to temperature and CO_2_ (Anav et al., 2013). Neglecting plant acclimation could contribute to these problems.

We do not advocate addressing acclimation by the accretion of additional model components and parameters. Instead, we propose an alternative theory-driven model development strategy. This strategy is based on eco-evolutionary optimality theory (Franklin et al., 2020), where eco-evolutionary refers to the fact that plants adjust to environmental conditions on both ecological and evolutionary timescales. This theory has been tested at various spatial and temporal scales, showing remarkable skill in predicting observed natural patterns at leaf (Wright et al., 2003; Maire et al., 2012; Prentice et al., 2014; Wang et al., 2017; Smith et al., 2019; Wang et al., 2020), plant (Franklin et al, 2012; Farrior et al., 2013; Lavergne et al., 2020) and ecosystem (Franklin et al., 2014; Baskaran et al., 2017) levels. So far, the theory has been tested at weekly to monthly timesteps, i.e. at the timescales of acclimation. However, to apply this theory in LSMs it needs to be tested at sub-daily timesteps, thus including both instantaneous and acclimated timescales.

Here we apply an existing optimality-based model for gross primary production (GPP), the P model (Wang et al., 2017; Stocker et al., 2020), to evaluate the potential of combining the two timescales in a parsimonious way. We extend the model to include both the instantaneous and acclimated responses in the simulation of GPP at a sub-daily timestep. We test the model using GPP derived from eddy covariance flux-tower measurements from boreal, temperate and tropical forests and also at a temperate grassland site. Our work provides a proof-of-concept for including acclimated responses in a LSM framework.

## 2 Materials and Methods

### 2.1 The P model

The P model (Wang et al., 2017) is an optimality-based model of GPP driven by solar radiation, temperature, vapour pressure deficit (VPD), ambient CO_2_ and the fraction of absorbed photosynthetically active radiation (fAPAR), which is assumed to be related to leaf area index (LAI) by Beer’s law (Stocker et al., 2020; Figure 1). Although most applications of the P model have used satellite-derived fAPAR data as inputs (Stocker et al., 2020; Wang et al., 2017), the model has also been run by estimating LAI from predicted GPP (Qiao et al., 2020) thus making it possible to project future conditions. Here, however, we use satellite-derived fAPAR as a model input. The P model is based on the Farquhar et al. (1980) biochemical model of photosynthesis (FvCB), but incorporates additional eco-evolutionary optimality hypotheses which express the acclimation of plant photosynthetic capacities and stomatal behaviour to environmental changes: the coordination hypothesis (Maire et al., 2012) and the least-cost hypothesis (Prentice et al., 2014). The coordination hypothesis states that plants tend to optimize their performance by adjusting their photosynthetic capacities (*V*_cmax_ and *J*_max_, Table 1) to use all of the available light. This leads to the conclusion that *V*_cmax_ and *J*_max_ should be continually adjusted to environmental variations according to general rules that do not depend on PFTs. The least-cost hypothesis states that plants minimize the sum of carbon and water costs — in terms of the maintenance costs for transpiration and carboxylation capacities. This hypothesis leads to an optimal ratio of the leaf-internal to ambient CO_2_ partial pressure (*c*_i_*:c*_a,_ Table 1) mathematically similar to Medlyn et al.’s (2011) formula (Prentice et al., 2014) but not requiring any parameters to be specified per PFT.

**Table 1.** Definitions of photosynthetic parameters, rates and constants used in the P model.

| Symbol | Description | Unit or value | Reference |
| --- | --- | --- | --- |
| $V_{\text{cmax}}$ | Maximum rate of carboxylation (or maximum rate of Rubisco activity) | $\mu\text{mol CO}_2 \text{ m}^{-2} \text{ s}^{-1}$ | Wang et al., 2017 |
| $J_{\text{max}}$ | Maximum rate of electron transport | $\mu\text{mol electrons m}^{-2} \text{ s}^{-1}$ | Wang et al., 2017 |
| $\chi = c_i : c_a$ | Ratio of leaf-internal to ambient partial pressures of $\text{CO}_2$ | Unitless | Prentice et al., 2014; Wang et al., 2017 |
| $c_i$ | Leaf-internal $\text{CO}_2$ partial pressure | Pa | Wang et al., 2017 |
| $c_a$ | Ambient $\text{CO}_2$ partial pressure | Pa | |
| $\xi$ | Sensitivity of $\chi$ to vapour pressure deficit (VPD or $D$) | $\text{Pa}^{1/2}$ | Wang et al., 2017 |
| $g_s$ | Stomatal conductance to $\text{CO}_2$ | $\mu\text{mol CO}_2 \text{ m}^{-2} \text{ s}^{-1}$ | Medlyn et al., 2011; Prentice et al., 2014 |
| $\phi_0$ | Intrinsic quantum efficiency of photosynthesis | $\text{mol mol}^{-1}$ | Rogers et al., 2017, 2019 |
| $\phi_0(T)$ | Temperature dependence function of quantum efficiency | $\text{mol mol}^{-1}$ | Bernacchi et al., 2003 |
| $\beta$ | The ratio of cost factors for carboxylation and transpiration capacities at 25 °C | 146 (unitless) | Stocker et al., 2020 |
| $c^*$ | The cost factor for electron-transport capacity | 0.41 (unitless) | Wang et al., 2017 |
| $K_C$ | Michaelis-Menten constant for carboxylation | Pa | Farquhar et al., 1980; Bernacchi et al. 2001 |
| $K_O$ | Michaelis-Menten constant for oxygenation | Pa | Farquhar et al., 1980; Bernacchi et al., 2001 |
| $K$ | The effective Michaelis-Menten coefficient for Rubisco kinetics | Pa | Farquhar et al., 1980 |
| $\Gamma^*$ | Photorespiratory compensation point | Pa | Farquhar et al., 1980; Bernacchi et al. 2001 |
| $\eta^*$ | Temperature dependence of the viscosity of the water, relative to its value at 25°C | unitless | Huber et al., 2009 |
| $P_a(z)$ | Atmospheric pressure at given elevation ($z$) | Pa | Berberan-Santos et al., 1997 |
| $J$ | Rate of electron transport | $\mu\text{mol electrons m}^{-2} \text{ s}^{-1}$ | Smith, 1937 |
| $A_C$ | Rubisco-limited assimilation rate | $\mu\text{mol CO}_2 \text{ m}^{-2} \text{ s}^{-1}$ | Farquhar et al., 1980 |
| $A_J$ | Electron-transport limited assimilation rate | $\mu\text{mol CO}_2 \text{ m}^{-2} \text{ s}^{-1}$ | Farquhar et al., 1980 |
| $A$ | Assimilation rate | $\mu\text{mol CO}_2 \text{ m}^{-2} \text{ s}^{-1}$ | Farquhar et al., 1980 |

**Figure 1.**
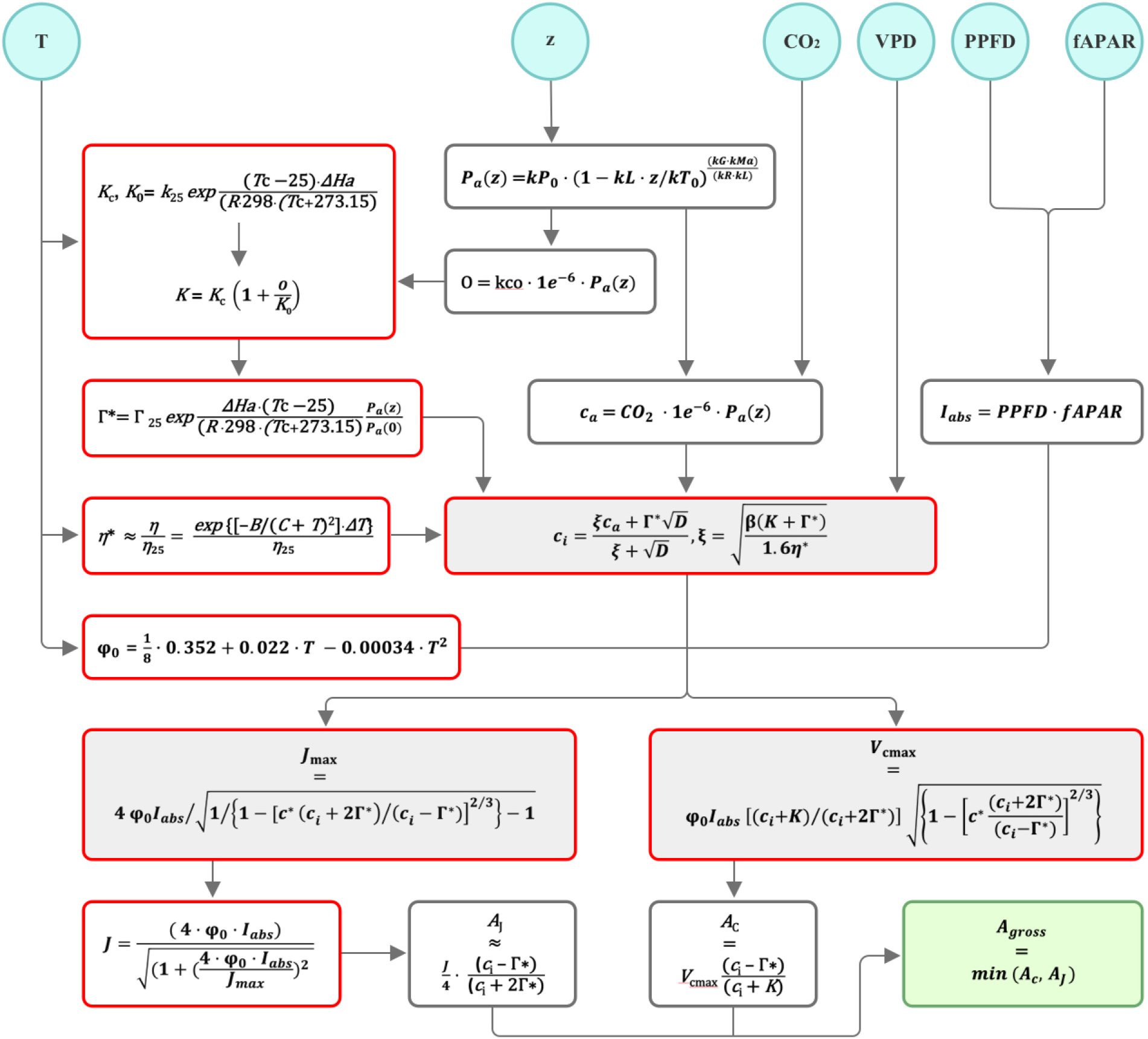
Flowchart of the P model. The light blue circles are model inputs, the red rectangles are the temperature dependent parameters, and the green rectangle is the model output. For details of the P model equations see Wang et al. (2017) and Stocker et al. (2020). For parameter definitions see Table 1 of this study.

The P model implicitly represents plant adaptation and acclimation, via photosynthetic capacity and stomatal behaviour, over a time scale of days and weeks. It reproduces observed variation in *V*_cmax_, *J*_max_ and stomatal conductance for CO_2_ (*g*_s:_ Table 1) along environmental gradients (Wang et al., 2017; Bloomfield et al., 2019; Smith et al., 2019; Dong et al., 2020; Wang et al, 2020). It also includes the measured effect of low temperatures on the intrinsic quantum efficiency of photosynthesis (φ_0_: Table 1) (Singsaas et al., 2001; Rogers et al., 2017, 2019). The parameters in the P model are either approximately constant and known from independent physiological studies, or estimated from analyses of independent data (*c*\* and β: see Table 1). The P model performs as well as more parameter-rich models (e.g. Zhang et al., 2019) at weekly to monthly time steps (Stocker et al., 2020), i.e. at the time scale of acclimation of key quantities such as *V*_cmax_ and *J*_max_.

### 2.2 Timescales of acclimation

To implement the P model at a sub-daily timestep requires an explicit distinction between the fast (instantaneous) response of photosynthetic rates (Fig. 1) and the slower acclimated response of photosynthetic traits (Fig. 1). To account for the acclimation of photosynthetic traits we use a running mean of the model inputs. We tested three approaches to find the optimal timescale for acclimation: the ‘daily’ approach computes a running mean of average daytime conditions; the ‘3 hours’ approach considers an average of three values from the middle of each day; the ‘noon’ approach considers only conditions around midday. The inputs are used to obtain the optimal values of *V*_cmax_ and *J*_max_ (eqs.1, 2). These represent the slow (acclimated) responses of the photosynthetic traits (Wang et al., 2017):

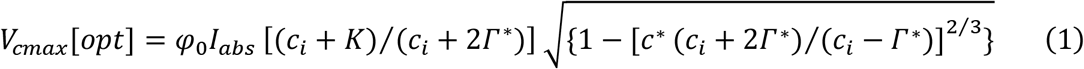

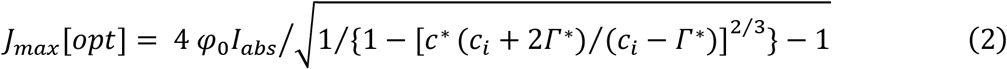

where *c*_i_ is the leaf-internal CO_2_ partial pressure (Pa), *Γ*\* is the photorespiratory compensation point (Pa), *K* is the effective Michaelis-Menten coefficient (Pa), *φ*_0_ is the intrinsic quantum efficiency of photosynthesis (mol mol^-1^), following the temperature dependence function *φ*_0_ (T) reported in Bernacchi et al. (2003), *I*_abs_ is the absorbed light, which is a product of the incoming photosynthetic photon flux density (PPFD, µmol photon m^-2^s^-1^) and fAPA. *c*\**=* 0.41 is a cost factor for electron transport capacity.

A standard function for the temperature response of *V*_cmax_ and *J*_max_, the Arrhenius equation (eq.3), is used to adjust both photosynthetic traits from the average to the actual temperature (eqs.3a, 3b). This adjustment for each half-hourly timestep represents the instantaneous response in the model.

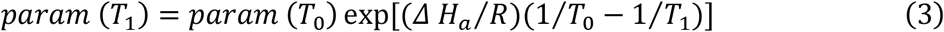

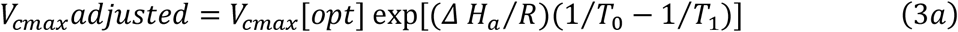

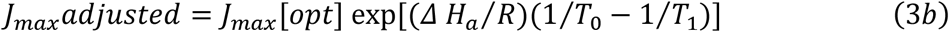

where T_0_ is the average temperature computed by a 15-day running mean (K) and T_1_ is the actual half-hourly temperature (K), *R* is the universal gas constant, and *ΔH*_a_ is the activation energy (*ΔH*_a_, J mol^−1^). The parameter values in the Arrhenius equation are summarized in Bernacchi et al. (2001, 2003).

Using the same logic for the stomatal conductance, we include a dynamic optimization of stomatal conductance operating on the *c*_i_*:c*_a_ ratio (χ, eq. 4) to obtain acclimated and optimal values of ξ, a parameter that determines the sensitivity of χ to VPD (Prentice et al., 2014). The acclimated response of ξ to environmental conditions (see e.g. Lin et al., 2015; Marchin et al., 2016) — is included in the *c*_i_ formula (eq.5), which is then adjusted to match the actual VPD to include the fast response of stomata to VPD:

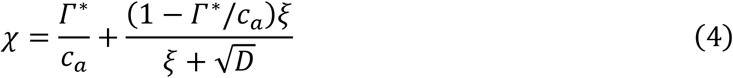

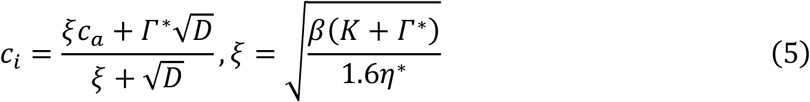

where *c*_i_ is the leaf-internal and *c*_a_ is the ambient CO_2_ partial pressure (Pa), *D* is the vapour pressure deficit (Pa), *β =* 146 is the ratio of the cost factors for carboxylation and transpiration capacities (at 25°C) (Stocker et al., 2020), and *η*\* is the viscosity of water relative to its value at 25°C.

As the flow chart illustrates (Fig.1), *V*_cmax_, *J*_max_, χ and thus *c*_i_, are necessary for computing the photosynthetic assimilation rates (*A*c and *A*_J_: Table 1). These are the two limiting rates for carbon assimilation; their minimum value gives the resulting GPP.

### 2.3. Incorporating acclimation in a land-surface modelling framework

Storing daily data in order to compute a running-mean would be computationally costly in a LSM context. We therefore tested whether the longer-term acclimation timescales could be mimicked based on a technique used to incorporate memory in other aspects of climate modelling, the exponential weighted moving average method. This method, here called the weighted mean approach, is used in forecasting systems that deal with inaccurate prediction caused by the insufficiency of historical observations and allows for a self-starting forecasting process without having to store past data (Yu et al., 2020). The method is used in a variety of applications in forecasting, from estimating soil moisture from precipitation (Campos de Oliveira et al., 2017) to vegetation acclimation processes (e.g. Vanderwel et al., 2015).

The weighted mean approach computes a mean in which the contribution of antecedent days decays exponentially with distance from the present. This is expressed by the exponential moving average (EMA) equation (see Supplementary Information, Text S1, eq.7). Together with the Arrhenius equation (see SI, Text S1, eq.8), these two formulae are used to update both photosynthetic traits (i.e. *V*_cmax_ and *J*_max_) in this new schema. The logic of the weighted mean differs from that of the running mean approach because, instead of operating on the inputs to the model it affects *V*_cmax_, *J*_max,_ and *c*_i_ directly. Specifically, the method uses the biochemical quantities at standard temperature, *V*_cmax25_ and *J*_max25_. This is because at standard temperature (25°C) *V*_cmax25_ reflects the quantity of active Rubisco in the canopy. First, the method requires the computation of optimal *V*_cmax_ and *J*_max_ (eqs.1, 2) based on conditions at noon; then, using the reciprocal formula of the Arrhenius equation (h^-1^; see SI, Text S1, eq.3a***) *V*_cmax25_ and *J*_max25_ are obtained (see SI, Text S1, eq.6) and used in the EMA equation. Computing the EMA equation, the acclimated responses of *V*_cmax,25_ and *J*_max25_ (for the current day) are obtained and then, with the canonical form of the Arrhenius equation (see SI, Text S1, eq.8), the instantaneous responses of both photosynthetic traits are computed at each half-hourly timestep. Like V_cmax,25_ and J_cmax,25_, ξ should vary slowly; however, there is no ‘fast’ reaction to temperature, so the Arrhenius function is not needed. After having obtained ξ for the current day, *c*_i_ is adjusted with the fast variation in VPD for each half-hourly timestep. Finally, these acclimated parameters — also adjusted to match the actual environmental conditions — are used to compute both photosynthetic rates (*Ac, A*_*J*_) and thus GPP at a sub-daily timestep.

To initialize the model simulations, we assume that on the very first day available in the dataset, the acclimated responses of *V*_cmax25_ or *J*_*max,25*_ (*V*_25,_ eq.7) are given by *V*_25,opt_ only. However the weight of *V*_25,opt_ decreases exponentially as time progresses (see SI, Text S1, eq.9). Therefore, it is necessary to apply a spin-up period of about 2 months before starting to look at the performance of the model. Then, we proceed with the application of eq.7 as discussed previously.

The EMA equation includes a parameter (α), the constant smoothing factor in time. According to eq.9 in SI (Text S1), α = 0.067 corresponds to about 15 days of memory. We therefore set α = 0.067 for consistency with the running mean method. We also tested a range of alternative values of α: 0.33, 0.143, 0.1, 0.067, 0.033, 0.022 and 0.0167, corresponding to 3, 7, 10, 15, 30, 45 and 60 days respectively.

### 2.4 Data and Evaluation

We compared model predictions with sub-daily observations from five sites in the FLUXNET2015 dataset (Pastorello et al., 2020) using the most recent common year (2014) for all five sites. We chose sites that represent a range of climate and vegetation types (Table 2): boreal forest (FI-Hyy), temperate deciduous broadleaf (US-UMB) and mixed (BE-Vie) forests, tropical forest (GF-Guy), and temperate grassland (CH-Cha). The FLUXNET data set provide meteorological variables (PPFD_IN, VPD_F, TA_F, CO2_F_MDS) on a half-hourly timestep at each site, as well as observed GPP. We used GPP based on the daytime partitioning method (GPP_DT_CUT_REF) (Lasslop et al., 2010; Pastorello et al., 2020). Since the FLUXNET2015 dataset does not provide fAPAR, we obtained this from the MCD15A3H Collection 6 dataset (Myneni et al., 2015). The MODIS FPAR product has a spatial resolution of 500 m and a temporal resolution of four days. We used the version of these data from Stocker et al. (2020) that has been filtered to remove data points where clouds were present and linearly interpolated from 4 days to daily. We used linear interpolation to derive fAPAR on the same sub-daily temporal resolution as the meteorological forcing.

**Table 2.** Summary of the characteristics of the FLUXNET2015 sites used for model evaluation.

| Site name | Site ID | Latitude (°) | Longitude (°) | Elevation (m) | Mean annual temperature (° C) | Mean annual precipitation (mm) | IGBP Vegetation type |
| --- | --- | --- | --- | --- | --- | --- | --- |
| Hyytiälä | FI-Hyy | 61.84741 | 24.29477 | 181 | 3.8 | 709 | Evergreen needleleaf forest |
| Vielsalm | BE-Vie | 50.30493 | 5.99812 | 493 | 7.8 | 1062 | Mixed forest |
| University of Michigan Biological Station | US-UMB | 45.5598 | -84.7138 | 234 | 5.83 | 803 | Deciduous broadleaf forest |
| French Guiana | GF-Guy | 5.27877 | -52.92486 | 48 | 25.7 | 3041 | Evergreen broadleaf forest |
| Chamau | CH-Cha | 47.21022 | 8.41044 | 393 | 9.5 | 1136 | Grassland |

We compared observed and simulated GPP over the growing season, where the growing season at each site was determined using the threshold approach defined by Lasslop et al. (2012). This approach defines the start and end of the growing season as the days that correspond to GPP values of >20% of the 0.05 and 0.95 quantile range.

The FLUXNET2015 data set provides information about the quality of data, through the quality-flag variables (see supplementary Table SM1 in Pastorello et al., 2020). We removed data points where the quality control (QC) is flagged as medium or poor-quality gap fill prior to comparison with model outputs. Times when there are no meteorological or GPP observations are necessarily ignored in the comparisons. There is no information that can be used to assess the quality of the fAPAR data.

Model goodness-of-fit was measured using *R*^2^, root-mean square error (RMSE) and the bias error (BE). The median RMSE, R^2^ and BE, were obtained by computing an RMSE, R^2^ and BE for each week during the growing season at each site. We estimated the number of weeks when model performance was reasonable by examining the RMSE distribution to determine a threshold to exclude outliers, which might be associated with data uncertainties.

## 3 Results

We tested the optimal timescale for acclimation to light availability by comparing simulations using average daily inputs, 3-hourly average inputs centred on noon, and midday conditions. The use of average daily inputs leads to a substantial underestimation of the observed GPP at all of the test sites. However, model predictions of GPP using 3-hourly or midday inputs are both consistent with the observations. At BE-Vie (Fig.2), for example, model performance using average daily inputs is poorer (R^2^: 0.92) than the either 3-hourly averages (R^2^: 0.98) or midday conditions (R^2^: 0.98). This is also the case at the other four sites (SI Figures S1, S2, S3, S4). These results support the hypothesis that plants coordinate their biochemical capacities to match the maximum level of light during a day, optimising to midday rather than average daytime conditions.

**Figure 2.**
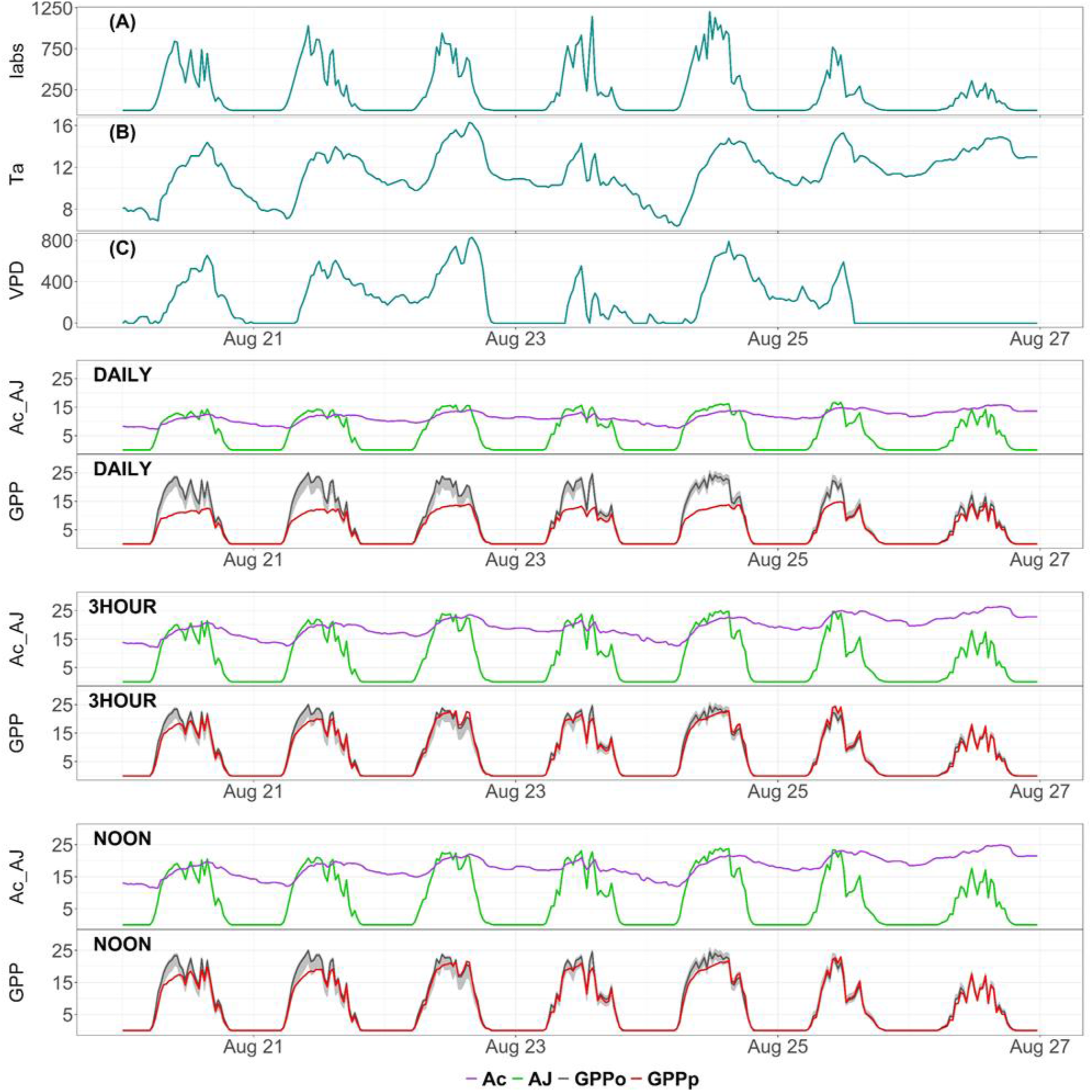
Sub-daily trends in model inputs (A, B, C) during one week in August 2014 at the Vielsalm (BE-Vie) site and the simulated Rubisco-limited assimilation rate (A_C_; µmol CO_2_ m^-2^ s^-1^), electron-transport limited assimilation rate (A_J_; µmol CO_2_ m^-2^ s^-1^) and gross primary production (GPP) using the running mean approach with inputs for average daytime conditions (DAILY), averaged over three hours from the middle of the day (3HOUR) and around midday (NOON). Simulated GPP (GPPp) is shown in red and the GPP derived from eddy covariance flux-tower measurements (GPPo) is shown in grey, both expressed in µmol CO_2_ m^-2^ s^-1^. Model inputs—Iabs, Ta, VPD—are in units of µmol Photon m^-2^ s^-1^, °C and Pa respectively.

Comparison of predicted and observed GPP at all five sites shows that the running-mean model accurately mimics diurnal cycles of GPP (Fig. 3). The median R^2^ over all weeks (Table 3) ranges from 0.88 (Ch-Cha, US-UMB) to 0.98 (GF-Guy). The median RMSE ranges from 3.89 µmol CO_2_ m^-2^ s^-1^ (Ch-Cha) to 2.28 µmol CO_2_ m^-2^ s^-1^ (BE-Vie). The median BE ranges from 2.04 µmol CO_2_ m^-2^ s^-1^ (GF-Guy) to 0.01 µmol CO_2_ m^-2^ s^-1^ (BE-Vie). There does not appear to be a relationship between the quality of the model fit and the length of the growing season. The model produces a good fit to observations at most of the sites for at least 80% of the individual weeks in 2014 (Table 3). The poorest performance in terms of number of weeks simulated accurately (68%) is for FI-Hyy and probably reflects uncertainties in the fAPAR inputs for this site.

**Table 3.** Summary of model performance statistics (RMSE is the root mean square error, R^2^ is the coefficient of determination and BE is the bias or systematic error). The number of weeks (No. weeks) is the length of the growing season at each site. The percentage of good weeks is estimated after excluding those weeks where the RMSE exceeds a threshold value of twice the median RMSE.

| Site ID | No. Weeks | Median RMSE | Median $R^2$ | Median BE | % Good Weeks |
| --- | --- | --- | --- | --- | --- |
| **BE-Vie** | 52 | 2.28 | 0.94 | 0.01 | 86.5 % |
| **FI-Hyy** | 38 | 2.53 | 0.91 | 0.99 | 68.4 % |
| **GF-Guy** | 52 | 3.67 | 0.98 | 2.04 | 82.7 % |
| **US-UMB** | 37 | 3.54 | 0.88 | 1.98 | 83.8 % |
| **CH-Cha** | 52 | 3.89 | 0.88 | 1.07 | 80.8 % |

**Figure 3.**
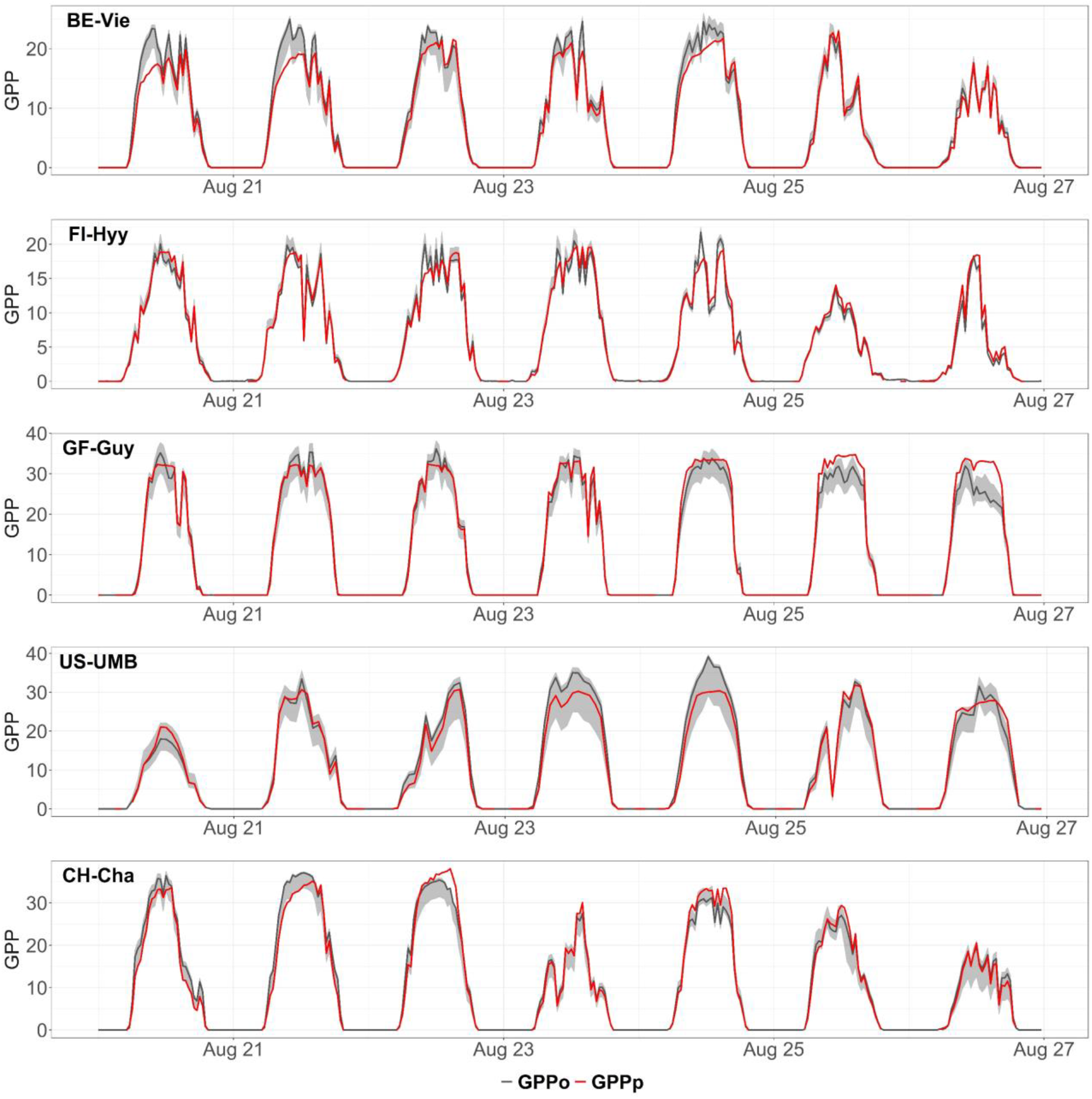
Comparison of simulated gross primary production (GPPp) and the GPP derived from eddy covariance flux-tower measurements (GPPo) for a single week in August 2014 at each of the five FLUXNET2015 sites (site IDs are displayed in the top left corner). GPP is expressed in µmol CO_2_ m^-2^ s^-1^.

**Figure 4.**
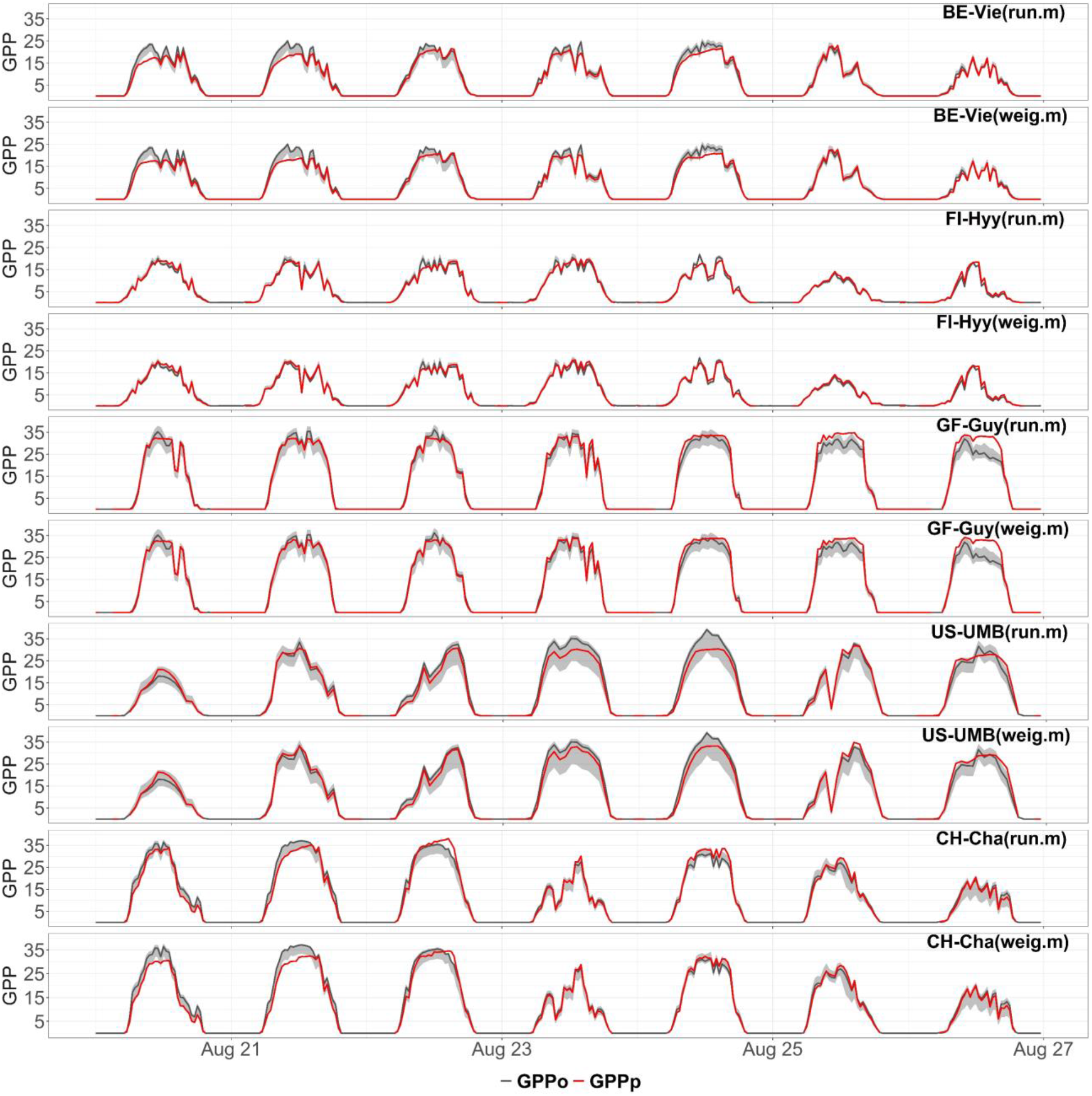
Comparison of the running mean (GPP_run.m) and weighted mean (GPP_weig.m) approaches for calculating gross primary production (GPP) for a single week in August 2014 at each of the five FLUXNET2015 sites. GPP is in units of µmol CO_2_ m^-2^ s^-1^.

The use of a 15-day period for calculating the running mean is a compromise. In three of the sites analysed, at shorter timescale (between three and seven days), there is the highest median RMSE and the lowest median R^2^ (SI, Fig.S5). After seven days the median R^2^ increases for each site, except for FI-Hyy, which also shows a sharper downward trend in the bias error than the other cases. Then, providing a longer timescale, beyond 15 days, there are no significant changes in the computed metrics, as the median RMSE and R^2^ are rather constant at four out of five sites. Although among these sites there are some differences, in most of the cases these results suggest that the optimum timeframe for acclimation is between 10 and 30 days, where 15 days is an average timeframe across all five sites (SI, Fig. S5).

Comparison between observed and simulated GPP shows very little difference between the running mean and weighted mean approaches. Visual comparison (Fig. 4) indicates that the two approaches produce essentially identical estimates of GPP at BE-Vie, FI-Hyy and GF-Guy. The weighted mean approach appears to produce marginally better results for the US-UMB site, but a marginally worse estimate at the CH-Cha site. Nevertheless, even for these two sites, the model performance using the weighted mean approach is consistent with the observed trends in the diurnal cycle. This shows it should be possible to include acclimation in LSMs at relatively low computational cost.

## 4 Discussion

We have developed a version of the P model that predicts GPP at sub-daily time scales. This model reproduces the diurnal cycle of GPP as recorded by flux-tower measurements across a range of different vegetation types. We have intentionally kept the model as simple as possible, as recommended e.g. by Prentice et al. (2015), in the interests of clarity. The model has few parameters, and their values are known from independent evidence. It does not distinguish between PFTs. We have succeeded in obtaining good simulations of flux data based on a minimal representation of the canopy as a big leaf. The model’s complexities (Fig. 1) are only those of the Farquhar et al (1980) model itself, and those necessary to implement optimality hypotheses that have been extensively tested – see e.g. Smith et al. (2019) for *V*_cmax_, Wang et al. (2017) for χ and the ratio *J*_max_:*V*_cmax_.

We have shown that plants adjust to midday conditions rather than average daytime conditions. It is reasonable to expect that plants would optimize to conditions during the midday period, when the light is greatest (Haxeltine and Prentice, 1996; Maire et al., 2012; Smith et al., 2019). We have also shown that the optimal timeframe for acclimation of carboxylation and electron-transport capacities, and the response of leaf-level carbon dioxide drawdown to vapour pressure deficit (VPD) is 15 days. Accounting for environmental variations over longer time periods does not produce significant differences in model performance metrics. In the original version of the P model (Wang et al., 2017; Stocker et al., 2020), designed to simulate GPP at weekly to monthly time steps, these acclimated responses are implicit. Our methodology makes it possible to separate the time scales of acclimation. Our analyses show that this distinction between fast and slow responses is essential to correctly predict plants’ responses to the environment.

The running mean and weighted averaging methods produce equally good simulations of the diurnal cycle of GPP. The weighted mean approach makes it possible to include acclimation in LSMs in a relatively straightforward way and to avoid prescribing different, fixed parameters for PFTs. Our results suggest that LSMs have had to specify different parameters for PFTs precisely because they do not represent acclimation. Plants growing in different environments have therefore had to be assigned different values of *V*_cmax25_ and *J*_max25_. But simulation would be more accurate, as well as requiring fewer parameters, if acclimation were allowed universally (in time and space) – so accounting more realistically both for seasonal variation, and for responses to environmental change.

## 5 Conclusions

We have adapted an existing optimality-based modelling framework to operate successfully at sub-daily timescale. The P model, without PFT-dependent photosynthetic parameters, accurately predicts GPP at half-hourly timestep across a range of different biomes. The method we propose is able to manage both timescales of acclimation. The weighted mean approach is suitable for implementation in a LSM. Our results suggest a way forward for LSMs to reduce their dependence on multiple parameters while, at the same time, taking into account plants’ acclimation to the environment.

## Acknowledgments, data availability and author contribution

GM and ICP acknowledge support from the ERC-funded project REALM (grant number 787203). SPH acknowledges the support from the ERC-funded project GC2.0 (Global Change 2.0: Unlocking the past for a clearer future, grant number 694481). GM acknowledges Gianpaolo Balsamo and Gabriele Arduini for useful discussions and technical assistance at the ECMWF centre. GM acknowledges Benjamin D. Stocker for providing fAPAR data.

The half-hourly implementation of the P model is generated with RStudio and is available through GitHub public repository: https://github.com/GiuliaMengoli/P-model_subDaily

Datasets for this research are available in these in-text data citation references: Pastorello et al. (2020), [Creative Commons (CC-BY 4.0) license], Stocker B. (2020, December 24), [http://doi.org/10.5281/zenodo.4392703]

GM, AAP, SB, SPH and ICP designed the study. ICP and GM developed the theoretical model. GM wrote the first version of the model code and conducted the analysis on the model outputs. CT implemented the current version of the model code and assisted in data preparation. AAP contributed to implementing the logic of the weighted mean approach. GM, ICP and SPH wrote the first draft of the manuscript. All the authors contributed to the final version of the paper.

