## Supplemental Text S1 for "Ecosystem photosynthesis in land-surface models: a first-principles approach"

[1] Department of Life Sciences, Imperial College London, Silwood Park Campus, Buckhurst Road, Ascot SL5 7PY, UK, [2] European Centre for Medium Range Weather Forecasts, Shinfield Road, Reading, RG2 9AX, UK, [3] Geography & Environmental Sciences, Reading University, Whiteknights, Reading, RG6 6AH, UK, [4] Division on Impacts on Agriculture, Forests and Ecosystem Services (IAFES), Fondazione Centro Euro-Mediterraneo sui Cambiamenti Climatici (CMCC), Viale Trieste 127, Viterbo, 01100, Italy, [5] Department of Biological Sciences, Macquarie University, North Ryde, NSW 2109, Australia, [6] Department of Earth System Science, Tsinghua University, Beijing 100084, China

### Contents of this file

Text S1

### Introduction

This supplementary information contains a more detailed explanation of the exponentially weighted moving average method.

**Text S1.**

**Description of the exponentially weighted moving average method**

The exponentially weighted moving average or exponential moving average (EMA) approach computes a mean in which the contribution of previous days decays exponentially with distance from the present. This is implemented by using the EMA equation (eq.7). Together with the Arrhenius equation (eq.8), the two formulae are used to update $V_{cmax}$ and $J_{max}$ in this new schema. While the running mean was computed on the model inputs, the weighted mean is applied to the photosynthetic traits directly. Specifically, $V_{cmax25}$ and $J_{max25}$, are the photosynthetic quantities used in this new schema. To follow the logic of this averaging method, here we report for simplicity only the $V_{cmax25}$ equations, since for $J_{max25}$ the formulae are the same, the only difference is that they include the optimal $J_{max}$.

The first step is to compute the optimal value of $V_{cmax}$ and $J_{max}$ based on current midday conditions (eqs.1,2), as illustrated in the running mean section (2.2). Then, their values at 25°C (here denoted as $V_{25}$) are calculated as follows:

$$V_{25,opt} = V_{cmax}[opt]\, h^{-1}(T_{noon}) \tag{6}$$

where $V_{cmax}[opt]$ is the optimal value of $V_{cmax}$ at noon obtained from eq.1, in which all parameters are based on conditions at the noon (half-hour) timestep; $h^{-1}(T)$ is the reciprocal of the exponential part of the Arrhenius eq.3 (see eq. 3a***); $T_{noon}$ is temperature (K) at the noon (half-hourly) timestep and consistent with the time at which $V_{25,opt}$ has been computed.

At this stage, the methodology requires the application of the EMA equation (eq.7) to obtain the new updated values of the two photosynthetic quantities, i.e. the acclimated responses of $V_{cmax25}$ and $J_{max25}$:

$$V_{25}[t] = \alpha\, V_{25,opt}[t] + (1 - \alpha)\; V_{25}[t-1] \tag{7}$$

where the acclimated value of $V_{cmax25}$ or $J_{max25}$ (denoted as $V_{25}$) is given by its optimal value at noon ($V_{25,opt}$) and its acclimated value from the previous day, weighted by α and (1- α) respectively. The smoothing parameter $\alpha$ is set here to 0.067 which corresponds to a memory of 15 days as explained in the *smoothing parameter* section below.

The acclimated responses of $V_{cmax25}$ and $J_{max25}$ are then adjusted to match with the actual temperature of the current day. Both quantities are updated at noon conditions by using the canonical form of the Arrhenius equation (h (T), eq.3a**), which allows to compute the instantaneous responses of both photosynthetic traits (here indicated as $V$) at each half hourly timestep:

$$V[t] = V_{25}[t]\, h\,(T) \tag{8}$$

where $V$ is the instantaneous value of $V_{cmax}$ or $J_{max25}$ at a given (half-hour) timestep. This will be used in the calculation of $A_c$ (and $A_J$) (Figure 1).

For $J_{max}$ the procedure is similar to $V_{cmax}$. Like $V_{cmax25}$ and $J_{max25}$, ξ should vary slowly; however, there is no 'fast' reaction to temperature, so Arrhenius function is not needed. After having obtained ξ for the current day, $c_i$ is adjusted with the fast variation in VPD for each half-hourly timestep. Then, these acclimated parameters, which are also adjusted to match the actual environmental conditions, are used to compute both photosynthetic rates ($A_c$, $A_J$) and thus the GPP at sub-daily timestep.

***Smoothing parameter (α)***

The smoothing-in-time coefficient α is an indicator of the length of the recent period to consider for the acclimation. This parameter ranges between 0 and 1 and expresses the contribution that past data (N-day memory) have on current values of photosynthetic traits, as illustrated here below.

Eq.7 can be re-written as the weighted average from the beginning of the simulation to current time *t*:

$$V_{25}[t] = \sum_{n=0}^{t} w[t-n]\ V_{25,opt}[t-n]; \quad w[t-n] = \alpha(1-\alpha)^n \tag{9}$$

where n is the index of the time steps (in days) back in time within the exponentially weighted moving average. The weight of $V_{25,opt}$ decreases exponentially as time progresses (e.g. weight of 20 days ago is $w(t\text{-}20)/w(t) = (1\text{-}a)^{20}$ smaller than at the current time). We can consider the e-folding time of the weights as the memory of the acclimation response. An e-folding decrease of the weights within a window of N days would imply a change in the weights of $(1\text{-}a)^N = 1/e$ which is equivalent to $log([1\text{-}a]^N) = -1$. Using the logarithmic power rule, we can derive $N = -1/log(1\text{-}a)$. For small $a$, $log(1\text{-}a)$ can be approximated by $-a$ (using Taylor expansion). Based on this, $N=1/a$ and inversely, $a=1/N$. Thus, a memory of 15 days (N=15) corresponds to $a$=0.067.

For consistency, having chosen a 15-day period of acclimation in the running mean method, we set α equal to 0.067. We also tested a range of alternative values of α: 0.33, 0.143, 0.1, 0.067, 0.033, 0.022 and 0.0167, corresponding to 3, 7, 10, 15, 30, 45 and 60 days respectively (Figure S5).

To initialize the model simulations, we assume that on the very first day available in the dataset, the acclimated responses of $V_{cmax25}$ or $J_{max25}$ ($V_{25}$, eq.7) are given by $V_{25,opt}$ only. However, as we mentioned above, the weight of $V_{25,opt}$ decreases exponentially as time progresses. Therefore, it is necessary to account for a spin-up period of about 2 months before starting to look at the performance of the model. Then, we proceed with the application of eq.7 as discussed previously.

***Arrhenius formulae*:**

$$param\ (T) = param\ \left(T_{ref}\right) \exp[(\varDelta\ H_a/R)(1/T_{ref} - 1/T)] \qquad (3)$$

where T is the temperature (K) and $T_{ref}$, by convention, is taken to be 298.15 K.

We can also write the same equation as:

$$param\ (T) = param\ \left(T_{ref}\right)\ h(T) \qquad (3a\ *)$$

where:

$$h\ (T) = \exp[(\varDelta\ H_a/R)(1/T_{ref} - 1/T)] \qquad (3a\ **)$$

And its reciprocal is:

$$h^{-1}\ (T) = \exp[(\varDelta\ H_a/R)(1/T - 1/T_{ref})] \qquad (3a\ ***)$$
