## Supplemental Figures S1 to S5 for "Ecosystem photosynthesis in land-surface models: a first-principles approach"

[1] Department of Life Sciences, Imperial College London, Silwood Park Campus, Buckhurst Road, Ascot SL5 7PY, UK, [2] European Centre for Medium Range Weather Forecasts, Shinfield Road, Reading, RG2 9AX, UK, [3] Geography & Environmental Sciences, Reading University, Whiteknights, Reading, RG6 6AH, UK, [4] Division on Impacts on Agriculture, Forests and Ecosystem Services (IAFES), Fondazione Centro Euro-Mediterraneo sui Cambiamenti Climatici (CMCC), Viale Trieste 127, Viterbo, 01100, Italy, [5] Department of Biological Sciences, Macquarie University, North Ryde, NSW 2109, Australia, [6] Department of Earth System Science, Tsinghua University, Beijing 100084, China

### Contents of this file

Figures S1 to S5

### Introduction

This supplementary information contains additional figures for the other sites, which are referred to in the results section of the main manuscript

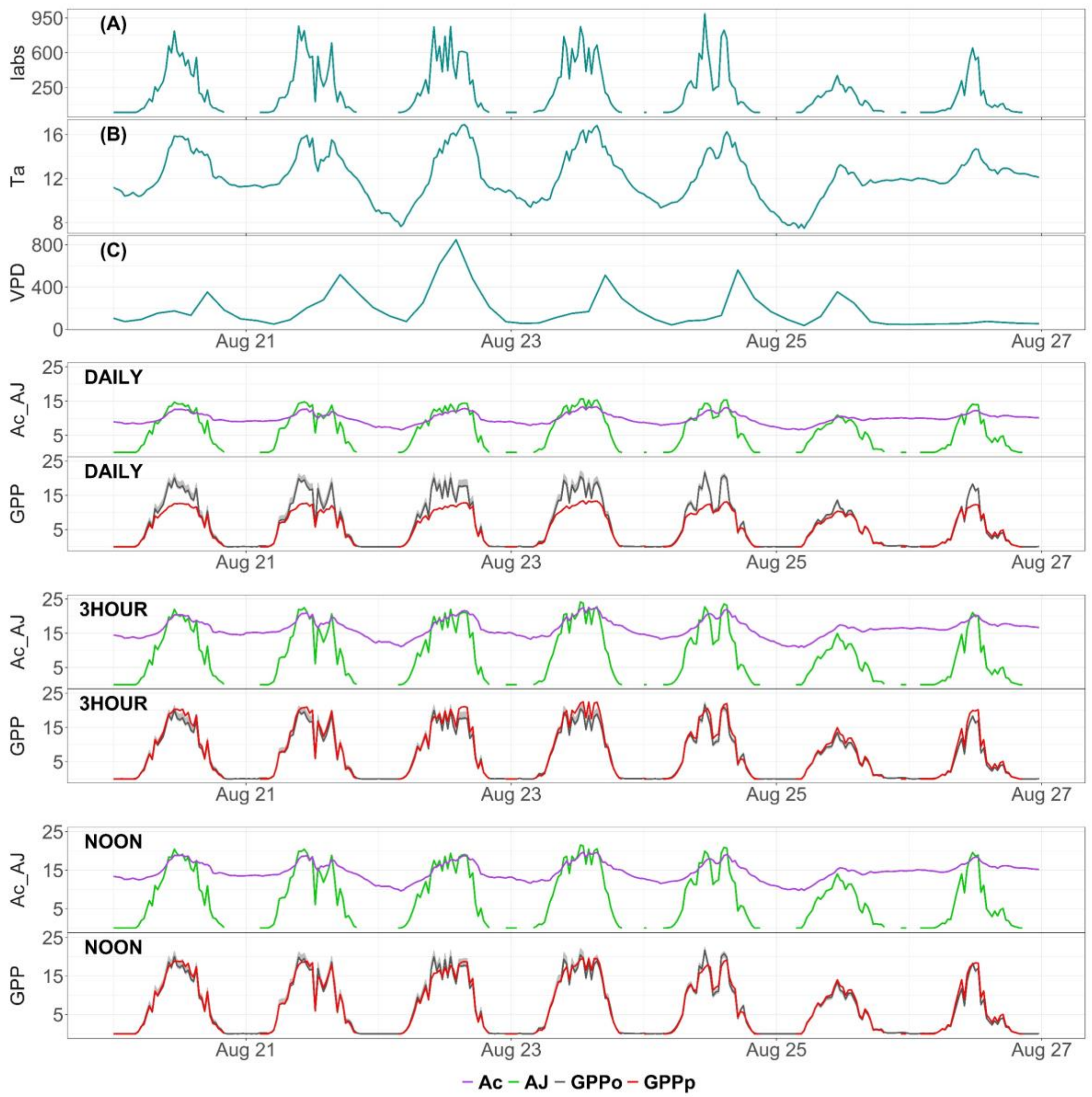


**Figure S1**. Sub-daily trends in model inputs (A, B, C) during one week in August 2014 at the **Hyytiälä (FI-Hyy)** site and the simulated Rubisco-limited assimilation rate ($A_C$; µmol $CO_2$ $m^{-2}$ $s^{-1}$), electron-transport limited assimilation rate ($A_J$; µmol $CO_2$ $m^{-2}$ $s^{-1}$) and gross primary production (GPP) using the running mean approach with inputs for average daytime conditions (DAILY), averaged over three hours from the middle of the day (3HOUR) and around midday (NOON). Simulated GPP (GPPp) is shown in red and the GPP derived from eddy covariance flux-tower measurements (GPPo) is shown in grey, both expressed in µmol $CO_2$ $m^{-2}$ $s^{-1}$. Model inputs—Iabs, Ta, VPD—are in units of µmol Photon $m^{-2}$ $s^{-1}$, °C and Pa respectively.

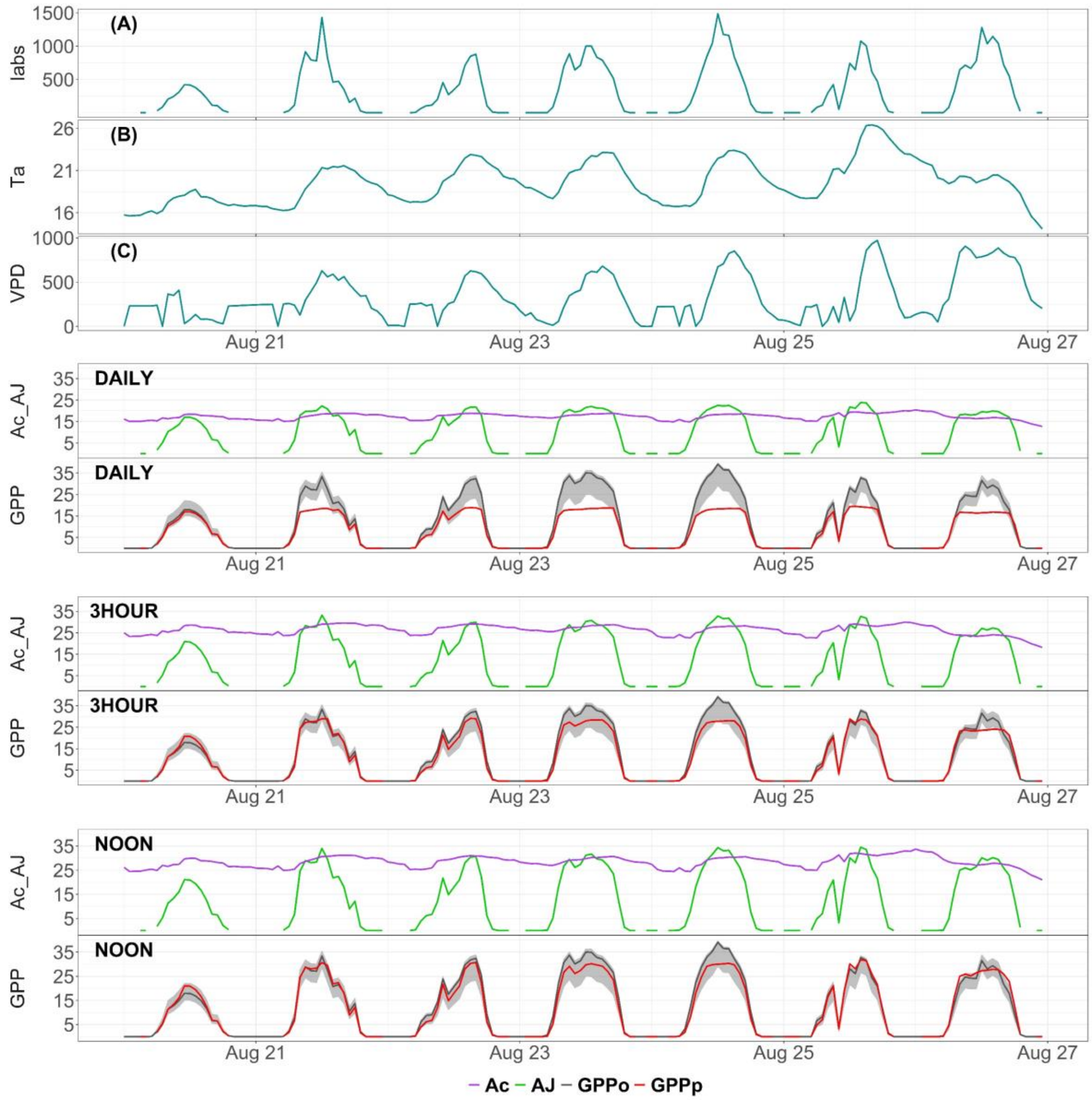


**Figure S2**. Sub-daily trends in model inputs (A, B, C) during one week in August 2014 at the **University of Michigan Biological Station (US-UMB)** site and the simulated Rubisco-limited assimilation rate ($A_C$; µmol $CO_2$ $m^{-2}$ $s^{-1}$), electron-transport limited assimilation rate ($A_J$; µmol $CO_2$ $m^{-2}$ $s^{-1}$) and gross primary production (GPP) using the running mean approach with inputs for average daytime conditions (DAILY), averaged over three hours from the middle of the day (3HOUR) and around midday (NOON). Simulated GPP (GPPp) is shown in red and the GPP derived from eddy covariance flux-tower measurements (GPPo) is shown in grey, both expressed in µmol $CO_2$ $m^{-2}$ $s^{-1}$. Model inputs—Iabs, Ta, VPD—are in units of µmol Photon $m^{-2}$ $s^{-1}$, °C and Pa respectively.

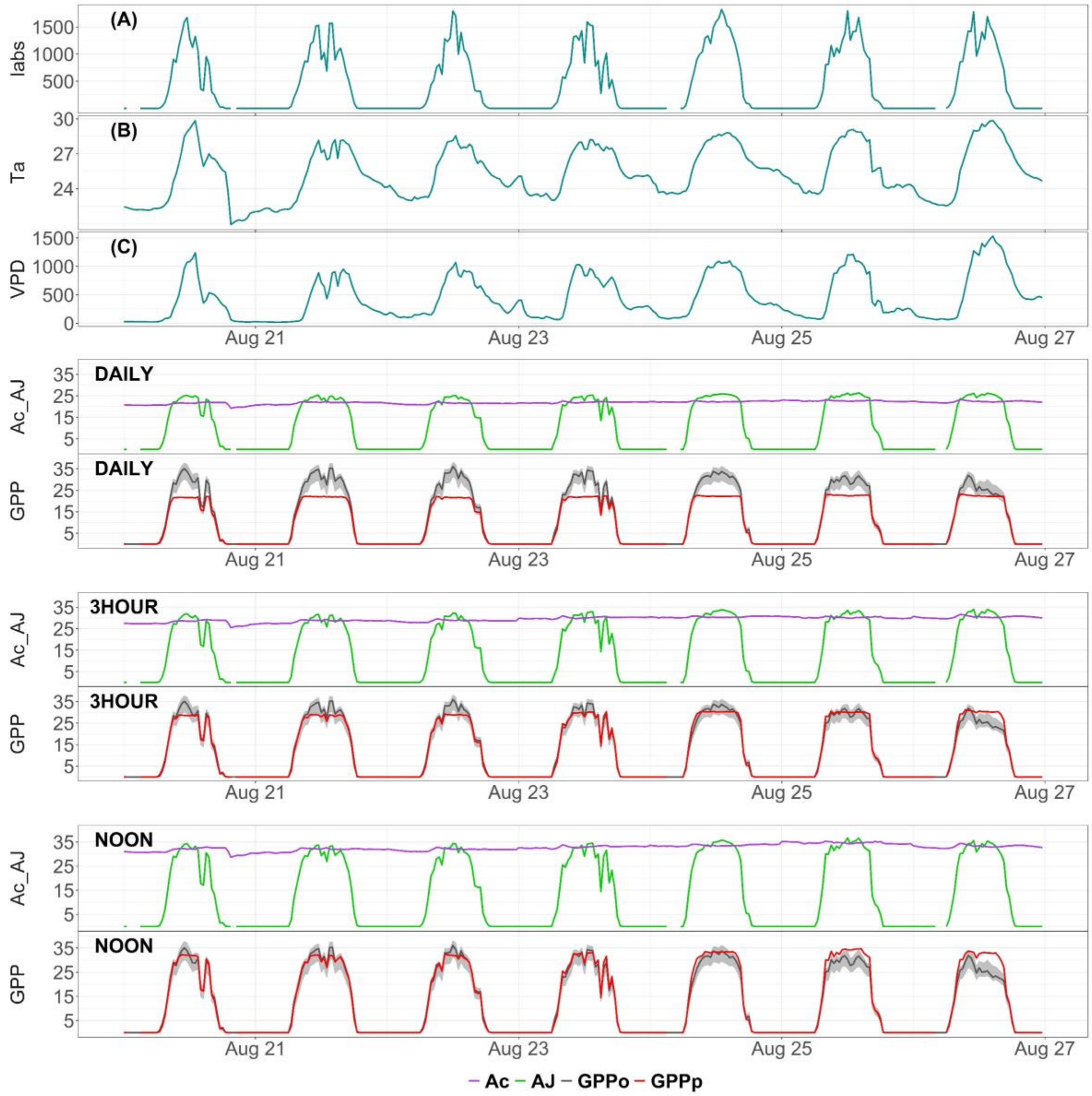


**Figure S3**. Sub-daily trends in model inputs (A, B, C) during one week in August 2014 at the **French Guiana (GF-Guy)** site and the simulated Rubisco-limited assimilation rate ($A_C$; μmol $CO_2$ $m^{-2}$ $s^{-1}$), electron-transport limited assimilation rate ($A_J$; μmol $CO_2$ $m^{-2}$ $s^{-1}$) and gross primary production (GPP) using the running mean approach with inputs for average daytime conditions (DAILY), averaged over three hours from the middle of the day (3HOUR) and around midday (NOON). Simulated GPP (GPPp) is shown in red and the GPP derived from eddy covariance flux-tower measurements (GPPo) is shown in grey, both expressed in μmol $CO_2$ $m^{-2}$ $s^{-1}$. Model inputs—Iabs, Ta, VPD—are in units of μmol Photon $m^{-2}$ $s^{-1}$, °C and Pa respectively.

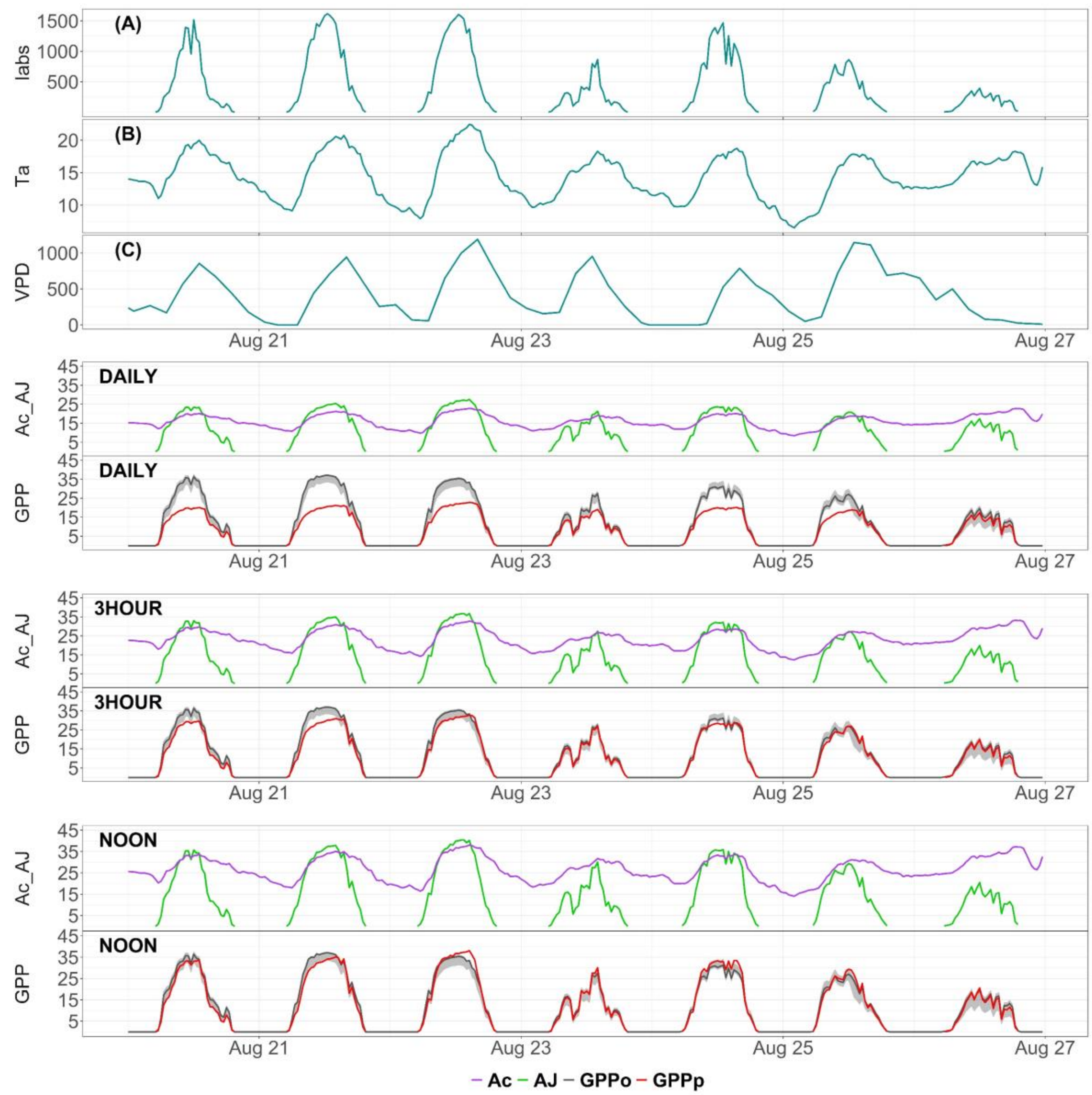


**Figure S4**. Sub-daily trends in model inputs (A, B, C) during one week in August 2014 at the **Chamau (CH-Cha)** site and the simulated Rubisco-limited assimilation rate ($A_C$; µmol $CO_2$ $m^{-2}$ $s^{-1}$), electron-transport limited assimilation rate ($A_J$; µmol $CO_2$ $m^{-2}$ $s^{-1}$) and gross primary production (GPP) using the running mean approach with inputs for average daytime conditions (DAILY), averaged over three hours from the middle of the day (3HOUR) and around midday (NOON). Simulated GPP (GPPp) is shown in red and the GPP derived from eddy covariance flux-tower measurements (GPPo) is shown in grey, both expressed in µmol $CO_2$ $m^{-2}$ $s^{-1}$. Model inputs—Iabs, Ta, VPD—are in units of µmol Photon $m^{-2}$ $s^{-1}$, °C and Pa respectively.

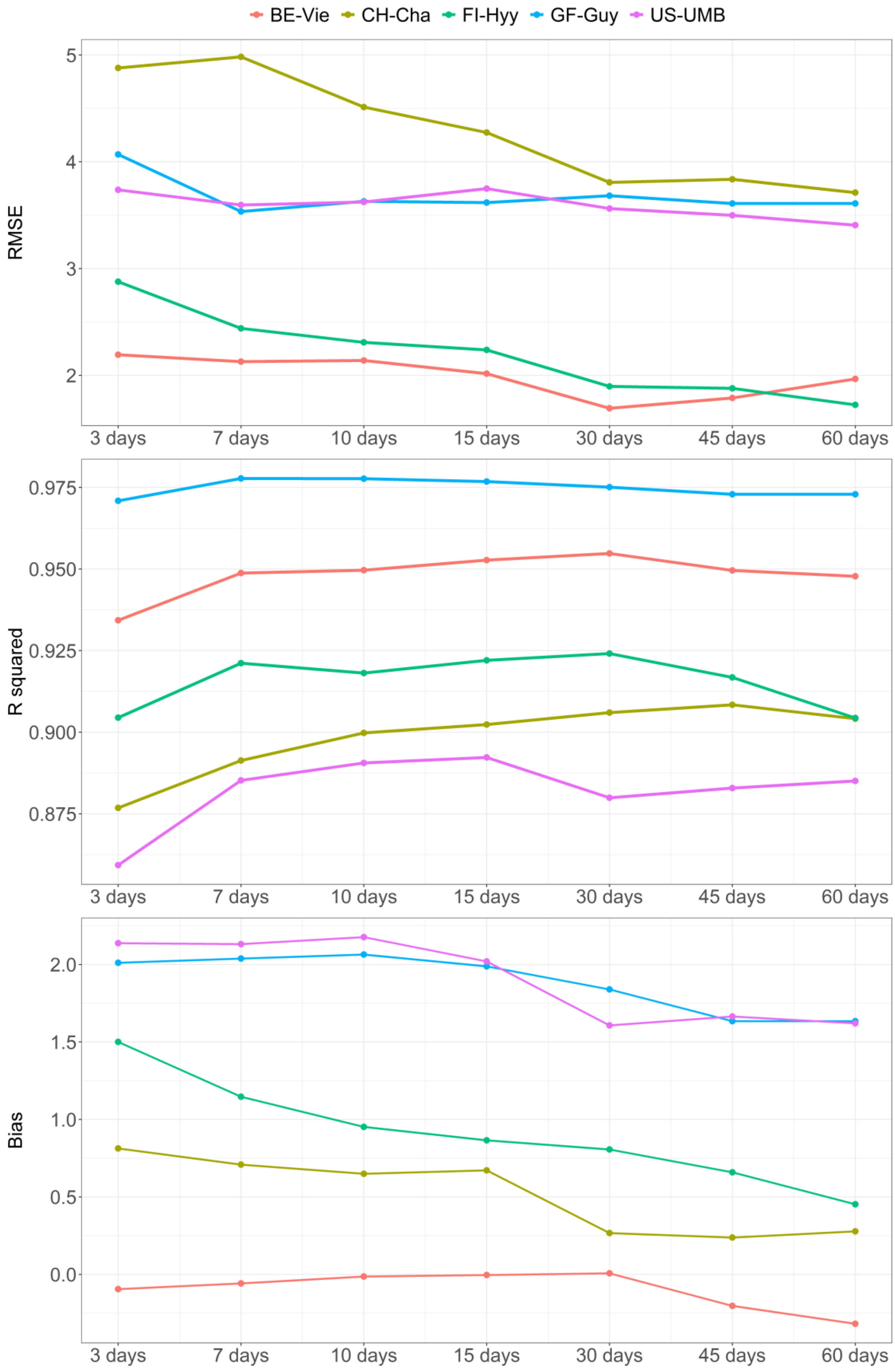

BE-Vie
CH-Cha
FI-Hyy
GF-Guy
US-UMB
RMSE
R squared
Bias
3 days
7 days
10 days
15 days
30 days
45 days
60 days

**Figure S5**. Median RMSE, $R^2$ and bias error distributions according to different lengths of acclimation time (n. of days) using the exponential moving average approach, computed per each of the five FLUXNET2015 sites, over 2014.
